# Agricultural land use and the sustainability of social-ecological systems

**DOI:** 10.1101/2020.07.27.222422

**Authors:** Diego Bengochea Paz, Kirsten Henderson, Michel Loreau

**Affiliations:** Centre for Biodiversity Theory and Modelling, Theoretical and Experimental Ecology Station, CNRS UMR 5321, 09200 Moulis, France

## Abstract

Agricultural land expansion and intensification, driven by human consumption of agricultural goods, are among the major threats to environmental degradation and biodiversity conservation. Land degradation can ultimately hamper agricultural production through a decrease in ecosystem services. Thus, designing viable land use strategies is a key sustainability challenge. We develop a model describing the coupled dynamics of human demography and landscape composition, while imposing a trade-off between agricultural expansion and intensification. We model land use strategies spanning from low-intensity agriculture and high land conversion rates per person to high-intensity agriculture and low land conversion rates per person; and explore their consequences on the long-term dynamics of the coupled human-land system. We seek to characterise the strategies’ viability in the long run; and understand the mechanisms that potentially lead to large-scale land degradation and population collapse due to resource scarcity. We show that the viability of land use strategies strongly depends on the land’s intrinsic recovery rate. We also find that socio-ecological collapses occur when agricultural intensification is not accompanied by a sufficient decrease in land conversion. Based on these findings we stress the dangers of naive land use planning and the importance of precautionary behaviour for land use management.

## 1 Introduction

Food production is the most basic and tangible example of humans’ dependence on nature. From Paleolithic hunter-gatherers, who relied on direct harvest from nature, to contemporary complex societies that rely on agriculture and livestock, human survival ultimately depends on what the land provides. An ever growing population and demand for food are putting unprecedented pressure on the environment (Tscharntke et al., 2012). Increased food consumption necessitates agriculture expansion; however, the last IPBES report (Bongaarts, 2019) highlights the role of agricultural land expansion as the main threat to biodiversity loss, mediated by the fragmentation and degradation of habitats (Corvalán et al., 2005; Jacobson et al., 2019; Nowosad and Stepinski, 2019). Degradation of the natural environment brings societal and economic consequences for human populations, as it can result in decreasing agricultural yields (Mitchell et al., 2014) and public health issues (Power, 2010). Conservation of biodiversity and natural spaces are often considered secondary objectives when compared to food security, but biodiversity and ecosystem services play an integral role in maintaining food supply. Agricultural productivity is strongly dependent on ecosystem services, such as pollination, nutrient cycling and pest control, that surrounding natural spaces provide (Mitchell et al., 2013). Therefore, conservation goals should not be seen as opposed to agricultural production or human well-being, as natural land is essential to provisioning services (Cazalis et al., 2018; Braat and de Groot, 2012). Allying natural and agricultural lands is the key to achieve sustainability and potentially avoid a socio-ecological collapse.

Throughout human history there have been numerous cases of societal collapse (Tainter, 1988). Examples of societal collapse are geographically diverse and have occurred over various time scales (Cumming and Peterson, 2017). They range from the Mesopotamian Akkadians in 2000 BC to the Easter Island society in 1300-1600 AD, in addition to the Western Roman Empire and the Lowland Mayans, in the Vth and IXth centuries of our era, respectively. This observation suggests that collapse is an intrinsic and common characteristic of human societies, rather than an exception. Despite the multitude of documented societal collapses, however there is little consensus on the mechanisms behind the decline of past civilisations.

The ecological point of view suggests that collapses can be explained by resource depletion, whether from over exploitation by the human population or as a result of environmental catastrophes (Tainter, 1988). The Eastern Islanders, the Anasazi, the Norse and the Mayans are typical examples of societies that are thought to have collapsed because of resource depletion. This ecological hypothesis, however, is still debated, as socio-political mechanisms are likely to have been involved. Nevertheless, it is unlikely that the collapse of a civilisation has a single cause. Mechanisms do not operate independently from each other, rather they act in synergy, as links of a causal chain (Tainter, 1988). For example, during the classic Mayan collapse, severe droughts decreased agricultural production, which led to increased warfare and political instability (Diamond, 2005). Environmental issues alter a society’s structure and may change human behaviours; in turn human actions and decision-making also shape the environment. A holistic understanding of societal collapse requires efforts to elucidate both the environmental and societal mechanisms behind it, as well as the feedbacks between them.

The empirical study of past societal collapses often relies on patchy archaeological data. Through modelling approaches, we are able to explore a variety of hypothetical scenarios and mechanisms, thus complementing classic empirical approaches. The early work of Brander and Taylor (1998) on the Easter Island collapse set in motion the use of socio-ecological models to address sustainability questions. Using a classical economics approach, they derived a model for the bi-directional coupling between the human population and its environment, similar to a prey-predator dynamical model, and demonstrated the potential role that resource over exploitation might have played in Easter Island’s societal collapse. Several others studies contributed to further exploring the Easter Island collapse, by expanding on Brander and Taylor’s initial model through the addition of new social and economic mechanisms (Brandt and Merico, 2015; Dalton and Coats, 2000; Good and Reuveny, 2006; Pezzey and Anderies, 2003; Reuveny and Decker, 2000). Both Ordinary Differential Equation and Agent Based Models have been proposed to quantify and understand past socio-ecological dynamics and collapses, in particular in the Anasazi (Axtell et al., 2002; Kohler et al., 2012), and Mayans(Heckbert, 2013; Kuil et al., 2016; Roman et al., 2018). Most studies have focused on societal mechanisms and climate variations (e.g., water scarcity) to explain the collapse. More recently, through a dynamical systems approach, (Roman et al., 2018) highlighted the crucial role that agricultural intensification may have played in the Mayans’ collapse, where demographic trends seemed consistent with increasing agriculture intensification.

The current environmental crisis has reignited the interest in societal collapse. There is a general agreement that overpopulation and overconsumption are the main threats to environmental conservation and sustainability (Barrett et al., 2020). Thus, recent studies have addressed sustainability questions by explicitly modelling human demography and consumption patterns. Theoretical and modelling studies are looking beyond the study of early societal collapses, in order to capture contemporary socio-economical mechanisms or derive general principles for a theory of collapse (Cumming and Peterson, 2017). Broadening the spectrum of possible connections between nature and human populations, Cazalis et al. (2018) built a model to explore socio-ecological dynamics through the dependence of humans on several ecosystem services. One study suggested that social inequalities and environmental degradation were present in all known past societal collapses (Motesharrei et al., 2014) and as such the authors proposed a general model showing various projections of socio-ecological collapses as a result of the feedbacks between social inequality and resource exploitation. More recently, Henderson and Loreau (2018) and Henderson and Loreau (2019) proposed a general theoretical framework to explain human demography across history in relation to resource accessibility, which can be used to explain the population explosion in the last century and potential future scenarios. Through an economic-ecological model Lafuite and Loreau (2017), Lafuite et al. (2017) and Lafuite et al. (2018) investigated how time lags in the response of biodiversity to anthropic perturbations can feedback on the human population via shortages in food production and undermine the sustainability of the socio-ecological system. These studies focus on food consumption as the link between humans and their environment, which is why we also focus on food production in this work.

The introduction of agriculture permitted the apparition of the first permanent human settlements. These Neolithic settlements were heavily reliant on the agricultural system and, as a result, when environmental disasters struck, the food supply and the population suffered (Downey et al., 2016). In some cases, as much as 60% of the population was lost due to failed crops. Over time technological developments made it possible for human societies to adopt more intense forms of agriculture, which increased resource production and food security. Agricultural production enabled the population to grow and allowed the development of complex societies via social differentiation and territorial expansion (Kuijt and Goring-Morris, 2002). This drive to increase agricultural production, however led to deforestation (DeFries et al., 2010), excessive freshwater use (Lilienfeld and Asmild, 2007), soil biodiversity loss (Tsiafouli et al., 2015), altered nutrient (Quinton et al., 2010) and water cycles (Davidson et al., 2012), decreased pollinator abundance, and increased vulnerability to environmental change, all of which can have deleterious effects on agricultural production. Agriculture is thus dependent on the natural environment, but it also heavily transforms this environment. Agriculture has been responsible for both the rise and fall of societies. The aim of future societies is to have agriculture improve social welfare, but how to achieve this is a major unknown.

Research on sustainable agricultural land use has led to the land sharing-sparing debate (Grau et al., 2013; Power, 2010). Whether it is better to protect larger areas of natural land and cultivate high-intensity fields on the remaining land; or protect smaller areas of natural land while practicing wild-life friendly, low-intensity agriculture, is a question that has not yet been fully answered. Different authors often arrive at different conclusions, some defending the sparing/intensification paradigm (Phalan et al., 2011a,b; Balmford et al., 2019) and others the sharing/agroecological one (Perfecto and Vandermeer, 2010; Power, 2010). The sparing-sharing debate has been criticized for omitting the coupling between land use and human demography (Phalan, 2018). Furthermore, the discussion generally examines discrete, opposing strategies, yet there is an entire spectrum between these two extremes. The impact of agricultural intensification on sustainability is an important issue at present, as in the developing countries foreign demand is fueling the conversion of large areas of natural land into intensively cultivated monocultures (Fearnside, 2001; Pengue, 2005; Reboratti, 2010; Soares-Filho et al., 2006). The result is a uniform landscape that is highly vulnerable to environmental fluctuations, destruction of natural habitats, fragmentation, contamination of underground water sources and nutrient runoff. These practices are detrimental to the environment, but agriculture is necessary to feed the population. It is obvious that a balance needs to be achieved between food production and natural land conservation, as the actions taken today could jeopardize the population’s viability in the long run.

Here we build a model to explore the effects of different agricultural land use strategies on long-term human-environment dynamics. Through a simple and tractable model accounting for the interaction between human demography and land dynamics, we study the viability of agricultural socio-ecological systems under different land use strategies along an intensification-expansion spectrum. We introduce a trade-off between intensification and the land conversion effort, and investigate for which land use strategies the population collapses due to land degradation. Our central premise, is that increasing agricultural production can promote further population growth. Thus, agricultural intensification, via increasing agricultural yields, can have a positive feedback on human demography, initiating the need for larger production and therefore causing further natural land conversion to agriculture, which eventually leads to a more degraded landscape. We test the conditions under which increasing agricultural intensification fails to spare enough natural land and promotes unsustainable population growth, pushing the environment through a tipping point and ultimately leading the social-ecological system to collapse.

## 2 Model description

### 2.1 Bidirectional coupling between human demography and land dynamics

Our model considers the conversion of natural land to agricultural land in relation to the demand from the human population. As population dynamics are driven by the resources humans can access and consume, they ultimately depend on the landscape’s composition. Resource production depends on the landscape composition but also on agricultural intensity. We conceive agricultural land use along two dimensions: the conversion effort, which controls the spatial extension of agricultural land, and agricultural intensity. In the model, humans adopt a land use strategy ranging from low intensity and high land conversion rates, to high intensity and low land conversion rates. This negative relation between agricultural intensity and the land conversion effort is grounded in the land sparing-sharing debate. Highly expansive and intense agricultural land uses have been identified as unsustainable. Hence the debate is whether the focus to achieve sustainability should be put on increasing intensification to reduce the converted areas, or extensification to have a wildlife friendly agricultural landscape. We aim to reproduce these two strategic poles by imposing a trade-off between agricultural intensity and land conversion effort, hence reducing the two strategical dimensions to a single parameter. In this study, we do not consider the evolution of the strategy over time, and assume it remains constant.

Agricultural land is exhausted and degraded, at different rates depending on the surrounding landscape, and ultimately becomes unproductive (Henderson and Loreau, 2019; Cramer et al., 2008). Natural land contributes to the recovery of surrounding land, acting, for example, as a species pool necessary for recolonization by native species (Cramer et al., 2008; Baeten et al., 2010). Hence, fragmentation of natural areas and degradation of natural patches surrounding degraded land can obstruct its spontaneous recovery. On the other hand, natural land can also become degraded. Indeed, a degraded state of land can propagate into a natural one, as is the case with a desertification front that propagates on semi-arid landscapes (Zelnik et al., 2017; Zelnik and Meron, 2018). The balance between the recovery and degradation processes depends not only on the extension of both natural and degraded land, but also on the borders between the two types of lands and on the level of degradation (Cramer et al., 2008).

### 2.2 Human demography

The human population *p* exhibits logistic growth (Suweis et al., 2013; Goldberg et al., 2016) with a carrying capacity that depends on resource production and consumption. We assume the carrying capacity to be the ratio between total resource production and per capita consumption *K*_*p*_ = *Y/C*, where *Y* is total resource production, depending on the landscape’s composition, and *C* is per capita resource consumption. For a given consumption intensity, the maximum number of humans that can be sustained is then given by the ratio of production over per capita consumption:

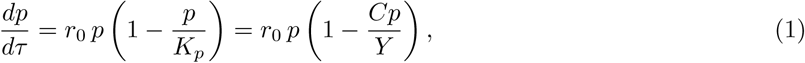

where *r*_0_ is the population’s growth rate at very low densities.

The amount of resources produced (*Y*) depends on the area of agricultural land (*a*), but also on that of natural (*n*) and degraded (*d*) land, as well as on agricultural intensity, *β*. Non-cultivated land, whether natural or degraded, provides ecosystem services that are crucial for agricultural production, such as pollination, nutrient cycling, pest control and water quality regulation (Mitchell et al., 2014). However, greater land degradation leads to fewer and lower-quality ecosystem services. Therefore, we do not consider natural and degraded land to contribute equally to agricultural production. Instead, we introduce an effective land function *E*_*l*_(*n, d, β*), which represents the effective area of non-cultivated land that provides ecosystem services to agricultural land:

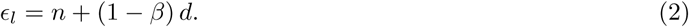

The contribution of degraded land to effective land decreases with its level of deterioration, which in turn depends on the level of agricultural intensification (*β*). We assume that more intensive agriculture results in higher degrees of land degradation and as such intensive agriculture transitions to highly degraded land. For simplicity, the contribution of degraded land to effective land decreases linearly with agricultural intensity.

We model agricultural resource production *Y* as the sum of the contributions from the total cultivated area (area contribution) and from the border of agricultural land with non-cultivated land, both natural and degraded, represented by the effective land *E*_*l*_ (border contribution). Therefore, the “area contribution” scales with agricultural land area and the “border contribution” with the square root of agricultural land area. Furthermore, we assume the relative weights of area and border contributions in production depend on agricultural intensity (*β*). As agricultural intensification grows, production, *Y*, becomes less dependent on the ecosystem services provided by the surrounding non-agricultural land and more dependent on human inputs. Hence, increasing intensification diminishes the border contribution and increases the area contribution on production. Therefore, we assume the area contribution increases linearly with agricultural intensity, *β*, while the border contribution decreases linearly with *β*.

The amplitude of the area and border contributions is modulated by the functions *Y*_*A*_(*β*) and *Y*_*B*_(*β*). These two functions can be interpreted as the characteristic productivity of the area and border contributions, respectively. For a given agricultural intensity (*β*), *Y*_*A*_(*β*) is the production per unit area of agricultural land and *Y*_*B*_(*β*) is the production per unit area of effective land per unit length of the agricultural land’s border.

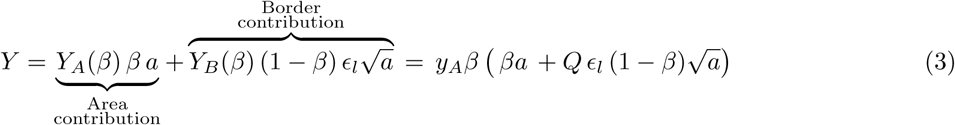

Agricultural intensity (*β*) ranges from 0 to 1, 0 being extreme low-intensity agriculture and 1 extreme high intensity. As intensification increases agricultural yields, we model the characteristic productivity of the area (*Y*_*A*_) and border (*Y*_*B*_) contributions as increasing functions of agricultural intensification. For simplicity, we assume a linear depdendency, i.e., *Y*_*A*_(*β*) = *y*_*A*_ *β* and *Y*_*B*_(*β*) = *y*_*B*_ *β*. The parameters *y*_*A*_ and *y*_*B*_ are then the productivities per unit of intensification. We introduce the parameter *Q* = *y*_*B*_*/y*_*A*_, which represents the relative importance of the border and area contribution to resource production.

Figure 2 shows the magnitude of agricultural resource production as a function of landscape composition. When *β* is close to 0, maximum production is obtained in a landscape where about a third of the land is agricultural. The food production is exclusively dependent on the services provided by the non-anthropogenic landscape. As we assume the services that non-cultivated land provides to agricultural land depend both on the area and quality of non-cultivated land and on the length of the border between them, at this extreme of the spectrum production scales with the square root of agricultural area. At the extreme, the fraction of natural land is not important because the degradation caused by the agricultural activity is extremely low, such that natural and degraded land contribute equally to effective land. As *β* grows, the fraction of natural land starts to have an impact, as degraded and natural land are not interchangeable anymore. When *β* approaches 1, production becomes exclusively dependent on agricultural land area. This is a scenario of extremely high agricultural intensity, where agricultural yields become independent of the services provided by the non-agricultural landscape, and rely exclusively on human inputs, such as fertilizers or pesticides. In the high intensity case, production is proportional to the area of agricultural land. Figure 2 also shows that yields increase with intensification, as maximum attainable production (yellow areas in the figure) grows with *β*.

**Figure 1:**
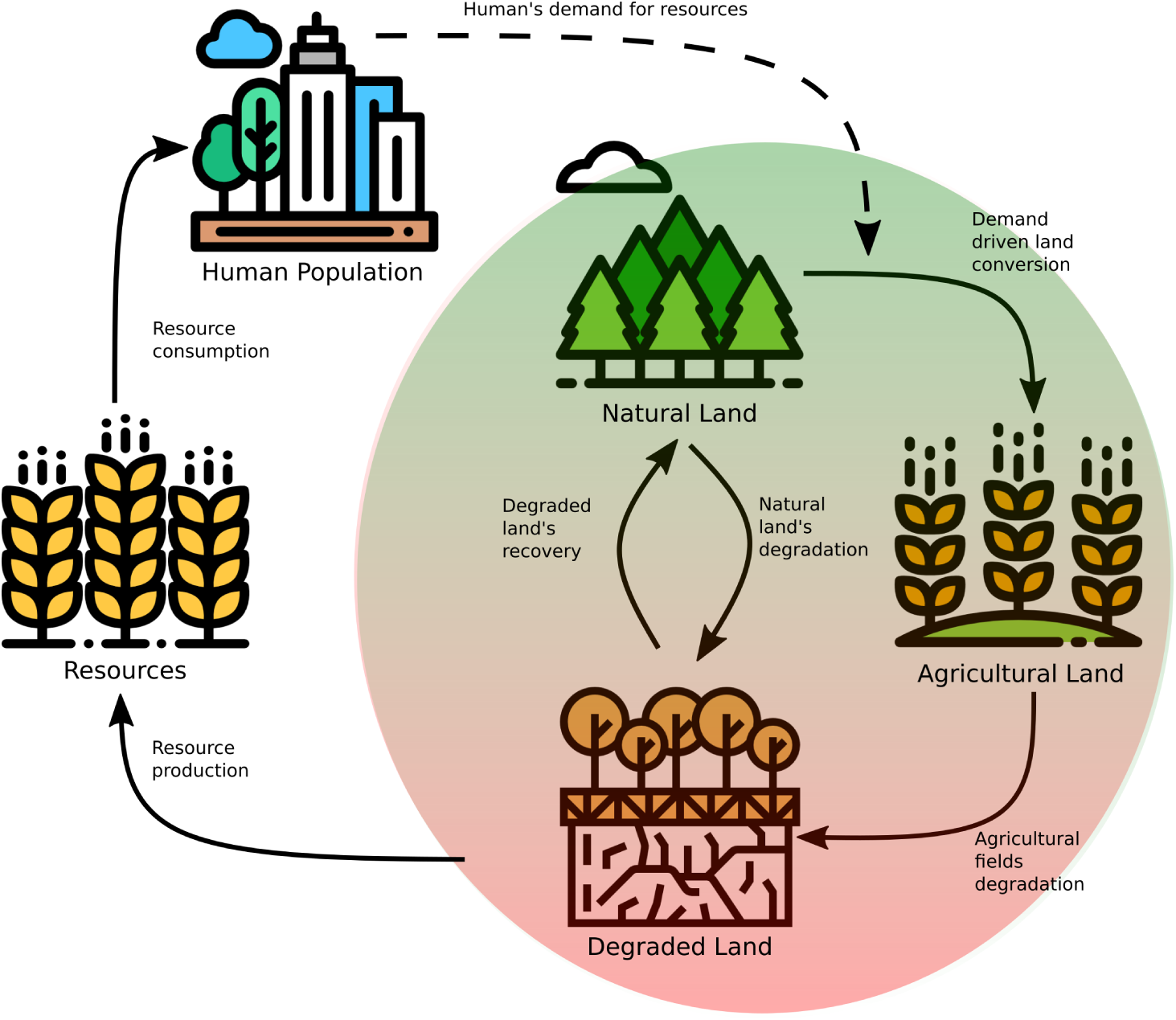
Model’s graphical representation. The right part of the diagram represents the landscape, composed of natural land, agricultural land and degraded land. The arrows between the three land types represent the possible land transformations we consider (conversion, recovery, degradation). Both the agricultural intensity and the landscape composition determine the amount of resources that are produced and consumed by the human population. Human demography is entirely determined by resource access. Changes in the human population size modify the population’s demand for resources and feedback on the landscape’s composition by increasing or decreasing the conversion of natural land for agricultural purposes.

**Figure 2:**
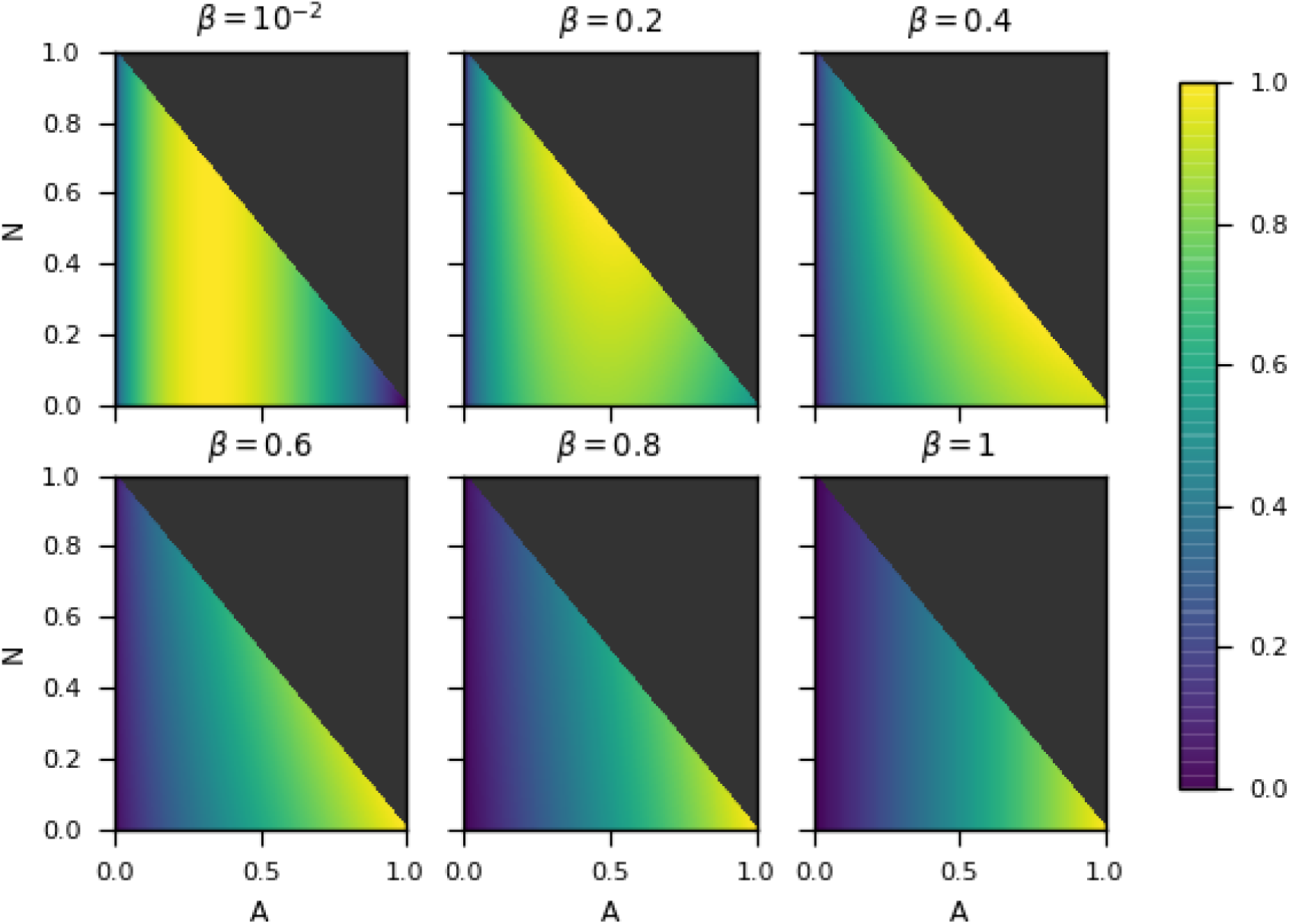
Agricultural production as a function of landscape composition for different agricultural intensities. The *x* and *y* axis correspond to the fraction of agricultural and natural land respectively. Each subplot corresponds to a different agricultural intensity (*β*). The production is normalised for each case, hence comparison of agricultural production’s magnitude between strategies is not possible. Instead, the figure shows the different effect that landscape composition has on each case. When agricultural intensity is very low (*β* = 10^*–*2^), production strongly depends on the services provided by the non-agricultural landscape, hence it decreases when the fraction of agricultural becomes bigger than a certain threshold (*A ≃* 0.4 in the plot). As intensification grows, natural land’s importance for production diminishes (*β* = 0.2, 0.4, 0.6, 0.8). On the high intensification extreme (*β* = 0.99), the production becomes exclusively dependent on human inputs, hence it grows monotonically with agricultural surface.

### 2.3 Land dynamics: agricultural land equation

Land conversion is driven by the human population’s demand for agricultural goods, which results on the conversion of natural land to agriculture. We assume that demand is equal to the total desired food consumption (*Cp*). Since we aim to investigate the impact of different land use strategies along the intensification-extensification spectrum on human-land dynamics, we impose a trade-off between agricultural intensity and land conversion rate. We model land conversion rate as a decreasing affine function of agricultural intensity (*β*). Nutrient runoff and soil erosion cause agricultural land degradation, which increases with intensity. Therefore we model the degradation rate of agricultural land as a linear function of *β*, capturing the fact that high-intensity agriculture degrades the land faster than does low intensity agriculture. The dynamical equation for the agricultural land area is given by

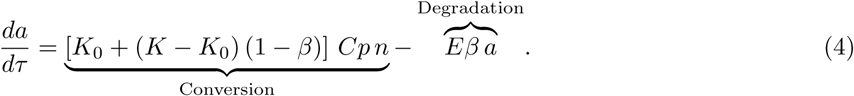

Parameters *K*_0_ and *K* are the minimum and maximum conversion rates per unit of demanded resources *Cp*, respectively. As the demand for resources is proportional to population density, *K*_0_ and *K* are also per capita rates of conversion. Therefore in the following we will call them per capita conversion rates. In the extreme high intensity scenario (*β* = 1), the conversion rate per person is at its minimum *K*_0_. In the extreme low intensity scenario (*β* = 0), the conversion rate per person is at its maximum *K*.

| Parameter name | Parameter description | Dimension | Non-dimensional | Value |
| --- | --- | --- | --- | --- |
| $r_0$ | Population growth rate at low densities | $\text{time}^{-1}$ | | |
| $y_A$ | Maximum production per cultivated area | mass/area | | |
| $C$ | Resources consumption per person | mass/person | | |
| $K_0$ | Minimum land conversion rate per resources demand | $\text{mass}^{-1}$<br>$\text{time}^{-1}$ | $k_0$ | 0.5 |
| $K$ | Maximum land conversion rate per resources demand | $\text{mass}^{-1}$<br>$\text{time}^{-1}$ | $k$ | 4.5 |
| $E$ | Maximum agricultural fields degradation rate | $\text{time}^{-1}$ | $e$ | 1.0 |
| $Q$ | Ratio between area and border contribution | $\text{length}^{-1}$ | $q$ | 1.0 |
| $R$ | Recovery rate of degraded land per length of frontier with natural land | $\text{length}^{-1}$<br>$\text{time}^{-1}$ | $r$ | [0.1 – 3.5] |
| $D$ | Maximum degradation rate of natural land per length of frontier with degraded land | $\text{length}^{-1}$<br>$\text{time}^{-1}$ | $d$ | [0.1 – 3.5] |
| $\beta$ | Sharing-sparing parameter | | $\beta$ | [0 – 1] |

### 2.4 Land dynamics: natural land equation

Apart from being converted to agriculture, natural land area can either increase through the spontaneous recovery of degraded land or decrease by the propagation of the degraded state of land. The natural land at the edges of degraded land fosters its spontaneous recovery through both biotic and abiotic processes. It acts as a species pool, promoting native species recolonization, or as a source of good quality water or chemical compounds to restore soil chemistry (Cramer et al., 2008; Baeten et al., 2010). The size of the natural patches is also important as larger patches foster more species and are more resilient to abiotic fluctuations (Mitchell et al., 2013, 2015). Hence, the recovery process depends both on the area of natural patches and on the size of their border with degraded land. We propose a spontaneous recovery term that scales both with natural land area *n* and with degraded land’s border 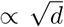 (Mitchell et al., 2015). The propagation of degraded land’s occurs throguh a symmetric mechanism, where the potential for degradation grows with degraded land area and with the natural land’s border. Therefore, the equation for the change in natural land is

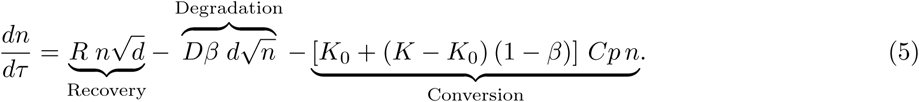

The parameters *R* and *Dβ* are the recovery and degradation rates, respectively. The degradation rate scales linearly with the agricultural intensification, such that more intensive agricultural land is more heavily degraded. Furthermore, heavily degraded land contributes to a greater extent to the degradation of natural land.

### 2.5 Non-dimensionalisation

We rescale the dynamical system by introducing the non-dimensional variables *P, N, A* and *t*, for population, natural land area, agricultural land area and time respectively:

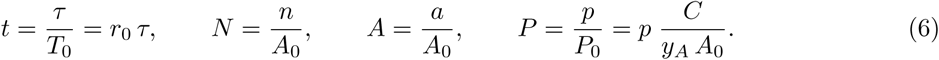

Time is rescaled to the characteristic timescale of human demography *T*_0_ = 1*/r*_0_. Parameter *A*_0_ is the total amount of land, hence variables *N* and *A* represent the fraction of natural and agricultural land in the landscape, respectively. We normalize the population by *P*_0_ = *y*_*A*_*A*_0_*/C. P*_0_ represents the population size that could be sustained if the whole landscape was cultivated with the highest intensity agriculture *β* = 1, given per capita consumption *C*. Indeed, when *β* = 1, if the whole landscape is cultivated, production is *Y* = *β*^2^ *y*_*A*_ *A*_0_ = *y*_*A*_ *A*_0_. The following dimensionless parameters emerge from the non-dimensionalisation: *k*_0_ – minimum land conversion rate, *k* – maximum land conversion rate, *e* – agricultural land degradation rate, *r* – spontaneous recovery rate of degraded land, *d* – degradation rate of natural land, and *q* – the relative importance of the border contribution to agricultural production.

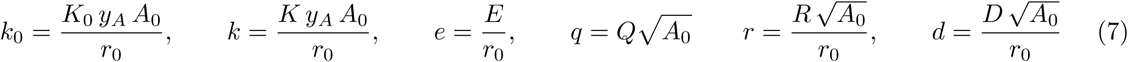

The dynamical equations describing the non-dimensional system behaviour are

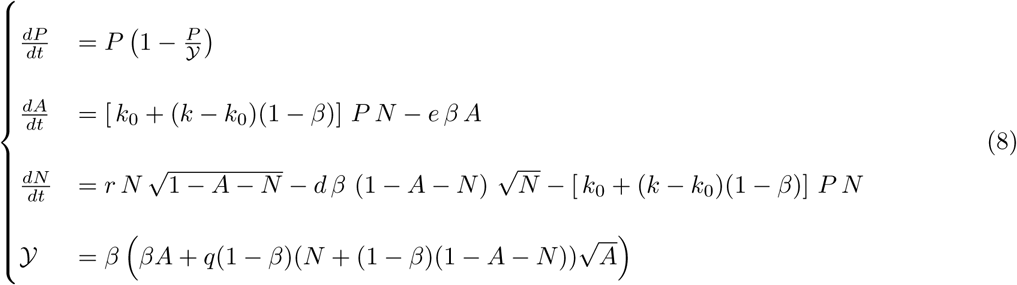

## 3 Results

### 3.1 Exploitation of a pristine landscape: sustainable vs. unsustainable land use strategies

We first look at the dynamics that follow the introduction of a small population in a pristine landscape. The time series are depicted in Figure 3. No matter the land use strategy, the early transient dynamics are identical. The human population converts the natural land into agricultural fields, thus increasing resource production, which positively feeds back on the human population. The increased population in turn accelerates land conversion. This positive feedback loop causes a population explosion accompanied by a transformation of the landscape. Agricultural land expansion fuels an increase in degraded land. Both land conversion and increasing amounts of degraded land contribute to the decline of natural land. The decrease in natural land area ultimately causes a deceleration of agricultural expansion until no more land is converted. The human population peaks with the agricultural area.

**Figure 3:**
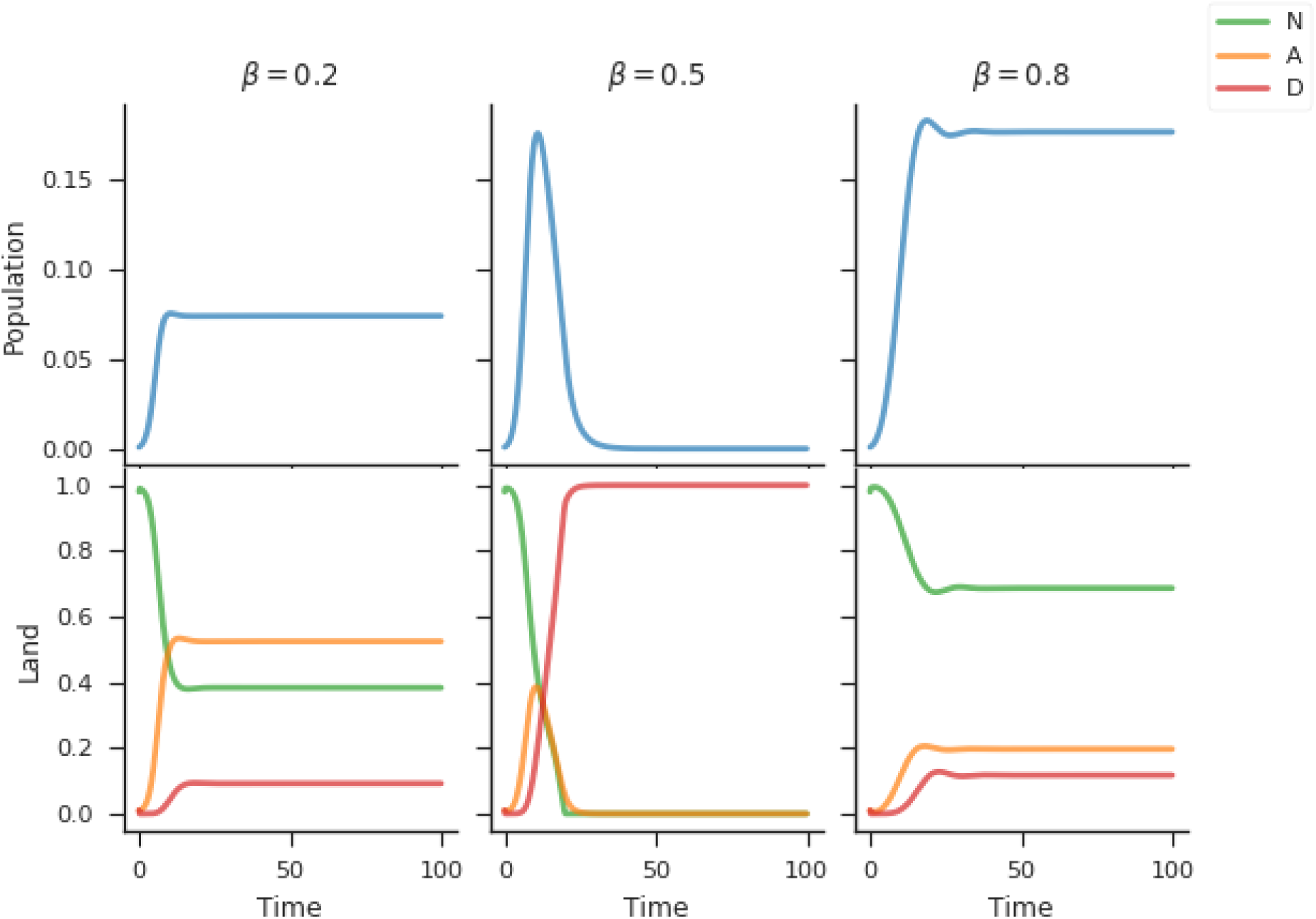
Temporal dynamics of the socio-ecological system for different agricultural land use strategies. *On the left:* dynamics emerging from low intensity agriculture and a high land conversion effort. The socioecological system reaches a viable equilibrium. *Center:* dynamics emerging from intermediate agricultural intensity and conversion effort. The socio-ecological system collapses. *On the right* : dynamics emerging from high intensity agriculture and low land conversion effort. A viable equilibrium is reached again. *Parameter values: r* = 1.0, *d* = 1.0, *k* = 4.5, *e* = 1.0, *k*_0_ = 0.5, *q* = 1.0.

Degraded land cannot be converted back to agricultural land. This introduces a time delayed feedback as the stock of natural land is not instantaneously regenerated. The time delayed feedback causesthe population to overshoot its carrying capacity. After the overshoot, the socio-ecological system can reach two different equilibria depending on the land use strategy *β*. We call *viable equilibrium* the one where the human population exists in the long term, and *collapse equilibrium* the one where the population goes extinct. In the viable equilibrium, the human population exists within a complex landscape, composed of a natural, agricultural and degraded land mosaïc. In contrast, the landscape in the collapse equilibrium is fully degraded. Without agriculture, there is no resource production and the human population cannot be maintained.

The land use strategy spectrum can be divided into three regions according to the system’s asymptotic behaviour, as a function of the strategy *β*. The first region corresponds to values of *β* between 0 and the transition to the collapse equilibrium at the critical point *β* = *β*_*c*,1_. We call this region the *sharing* side of the spectrum, as land use strategies in that range mimic land-sharing kinds of strategy (e.g. low intensity agriculture over large areas). The *collapse range* (referred to as Δ*β* later in the text) refers to the region of the spectrum where strategies lead to the collapse equilibrium. When land use strategies are inside the collapse range, the degraded land propagates into the whole landscape, leading to a population collapse. The region between the collapse range and the viable equilibrium is designated the sparing side of the spectrum, as the strategies in this region mimic land-sparing kind of strategies (e.g. high intensity agriculture over small areas).

### 3.2 On the path to socio-ecological collapse

The existence of the collapse range is due to changes in the stability of the viable equilibrium as a function of the agricultural land use strategy *β*. On the sharing side of the spectrum, the viable equilibrium is a stable focus-node. Hence, in the phase space, trajectories follow spirals before reaching the fixed point (Figure 4 (a)), which translate into damped oscillations over time (column (a) of Figure 5). As land use strategies come closer to the collapse range (*β* increases), the amplitude of the oscillations grow, which delays the system’s convergence to the viable equilibrium. When the land use strategy enters the collapse range, the viable equilibrium becomes a saddle-focus and loses stability (Figure 4 (b)). The stability loss is caused by a subcritical Hopf bifurcation which leaves the collapse equilibrium as the sole stable attractor for the socio-ecological system.

**Figure 4:**
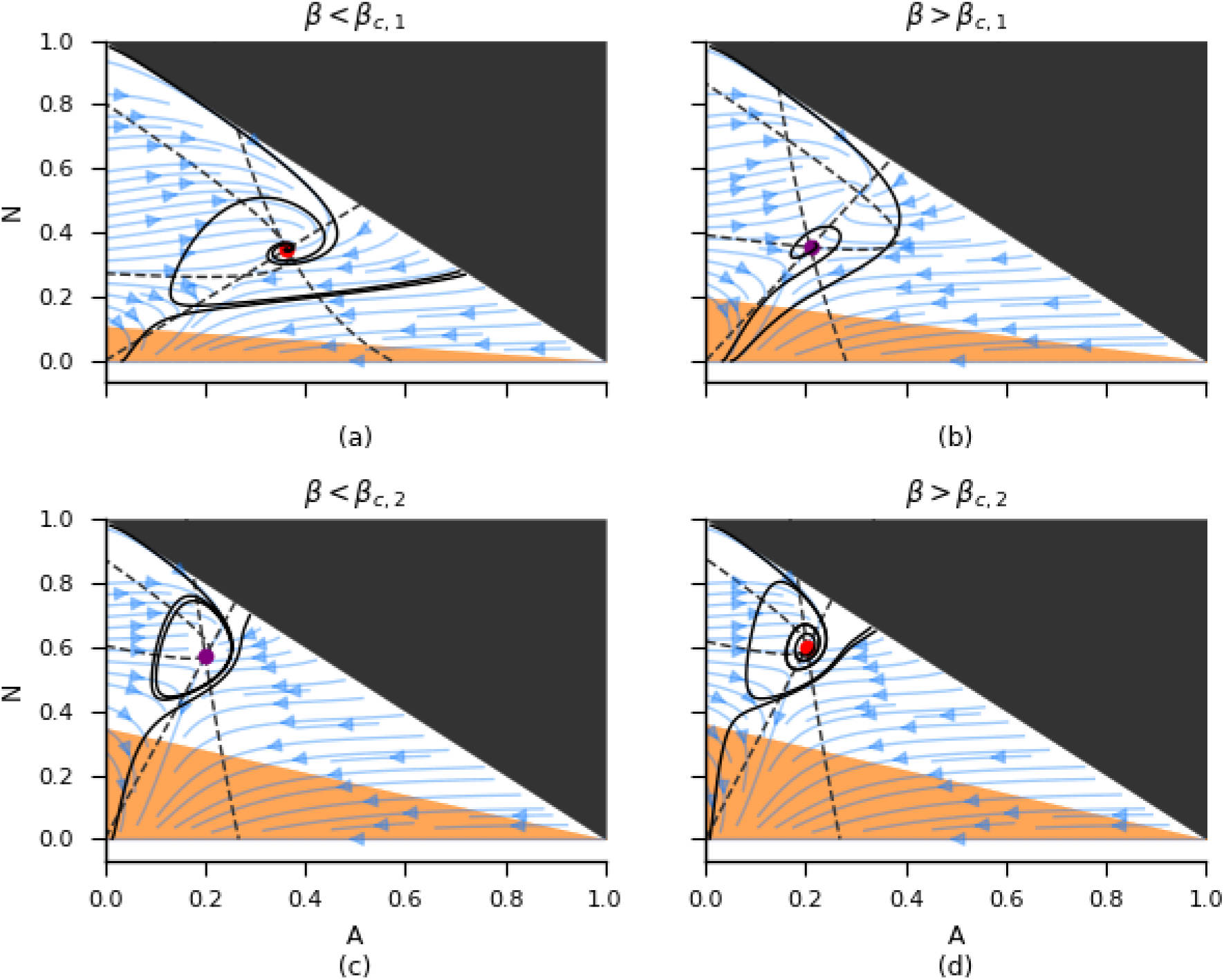
Phase representations of the socio-ecological system for strategies in the viable and collapse regions of the strategy spectrum. The plots correspond to particular landscape planes of the three dimensional phase space. The chosen planes are the ones containing the system’s viable equilibrium, hence determined by setting the population at its viable equilibrium value. The dotted grey lines are projections of the nullplanes for the population, the natural land and the agricultural land on the chosen landscape plane. The black solid lines are projections of simulated trajectories. The vector field depicted with blue arrows indicates the landscape’s direction of change for each landscape composition given a population at equilibrium. *On the top:* before (a) and after (b) the first transition to collapse. A subcritical Hopf bifurcation causes the stability loss of the viable equilibrium explaining the transition to collapse. *On the bottom:* emergence of a stable limit cycle (c) from a stable focus node (d) after a supercritical Hopf bifurcation. *Parameter values: r* = 1.0, *d* = 1.0, *k* = 4.5, *e* = 1.0, *k*_0_ = 0.5, *q* = 1.0.

**Figure 5:**
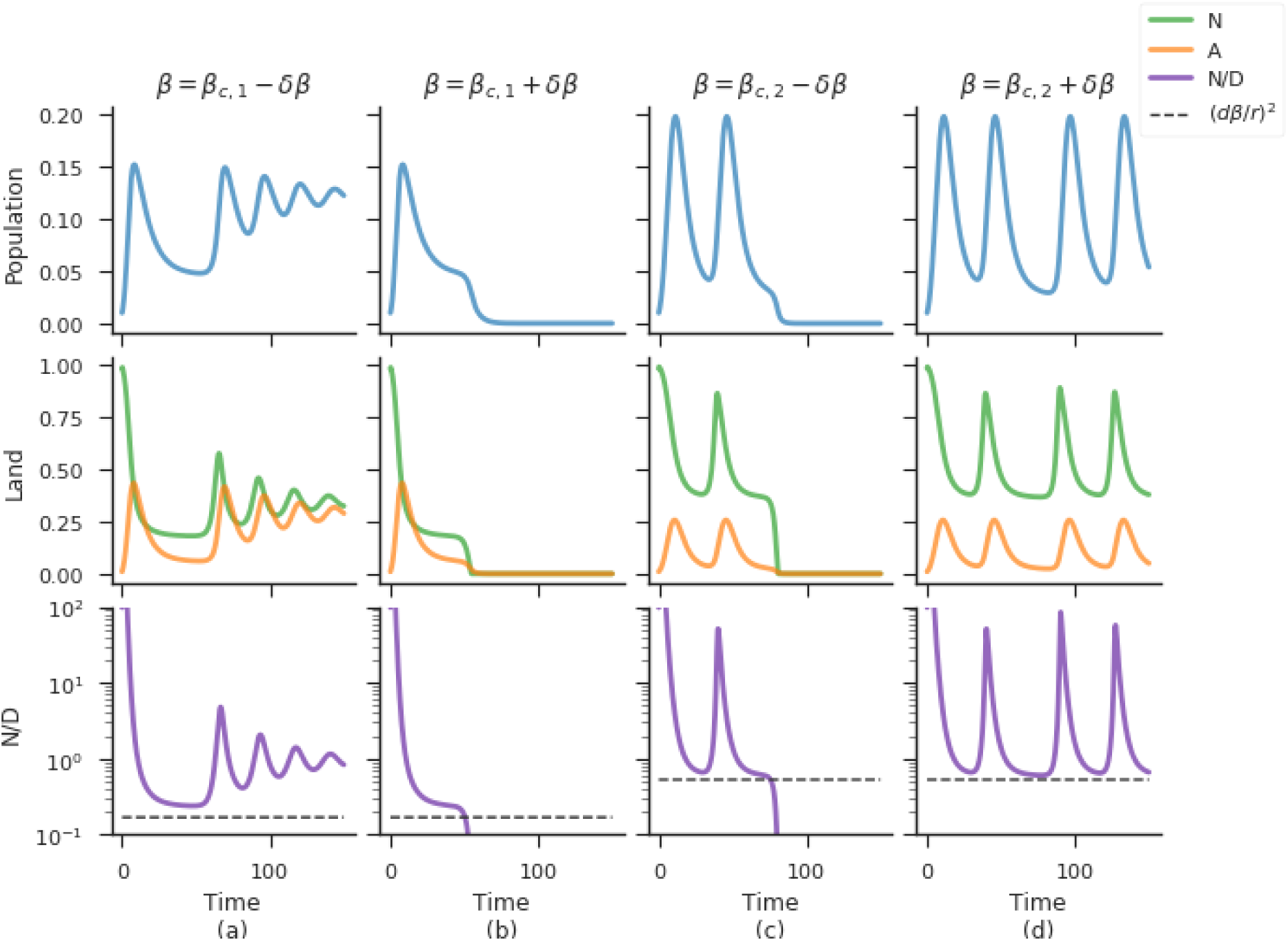
Temporal dynamics of the socio-ecological system at the edges of the collapse range. *Column (a):* dynamics before the first transition to collapse. The equilibrium is reached after large amplitude damped oscillations. *Column (b):* dynamics after the first transition to collapse. *Column (c):* dynamics after the second transition to collapse. The growth of the oscillations’ amplitude pushes the system through a threshold and causes the collapse (bottom plot). *Column (d):* dynamics before the second transition to collapse. The collapse is avoided as the system oscillates around the equilibrium without reaching it. *Parameter values: β*_*c*,1_ = 0.4106108, *β*_*c*,2_ = 0.7272030, *δβ* = 10^*–*7^, *r* = 1.0, *d* = 1.0, *k* = 4.5, *e* = 1.0, *k*_0_ = 0.5, *q* = 1.0.

On the sparing side of the spectrum, the transition to collapse has a different origin. As for the sharing side of the spectrum, the system converges to a viable equilibrium via damped oscillations (Figure 4 (a)) which grow in amplitude as the collapse range is approached. However, in this case the system undergoes a supercritical Hopf transition when the critical point is reached. Hence, the stability loss of the viable equilibrium is accompanied by the birth of a stable limit cycle, which allows the socio-ecological system to potentially escape the collapse equilibrium (Figure 4 (c)) and oscillate around the viable equilibrium. However, the amplitude of the oscillations grow as the land use strategy moves in the sharing direction (*β* decreases). Eventually, the oscillations become large enough to push the system through a tipping point provoking a socio-ecological collapse (bottom of column (c) in Figure 5).

Analytically, we can determine a threshold landscape composition after which socio-ecological collapse is unavoidable. Analysis of the natural land’s dynamical equation gives the following condition

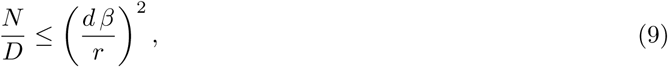

where *D* = 1 *– A – N* is the fraction of degraded land. The threshold depends on the land recovery potential *r*, as well as on the degradation potential *d β*. The threshold represents the point at which the landscape is so deteriorated that the remaining fraction of natural land is not sufficient to recover the degraded land nor to maintain its natural state. Hence, degraded land starts propagating into the natural land, resulting in the complete degradation of the landscape and population extinction. Close to the second transition to collapse, oscillations approach the previous threshold (column (d) of Figure 5). A small change in the land use strategy increases the oscillations’ amplitude and pushes the system through the tipping point, driving it to collapse (column (c) of Figure 5).

### 3.3 Role of land recovery potential on the size of the collapse range

The size of the collapse range Δ*β* = *β*_*c*,2_ *– β*_*c*,1_ is highly dependent on the degraded land’s recovery potential (*r*), as Figures 6 and 7 show. As the land recovery potential increases, the size of the collapse range decreases until it disappears. It is interesting to note that on the sharing side of the spectrum, when *β < β*_*c*,1_, an increase in *β* leads to higher agricultural yields and larger populations. However, the decrease in the land conversion effort is not high enough to prevent the fraction of natural land to decrease. Indeed, agricultural intensification increases the natural land’s degradation rate *βd*, increasing the potential of degraded land to propagate into the rest of the landscape. When agricultural intensification is not accompanied by a sufficiently large reduction in the conversion effort, the system enters the path to collapse. At the critical point, the levels of degradation are sufficiently high to cause a socio-ecological collapse.

**Figure 6:**
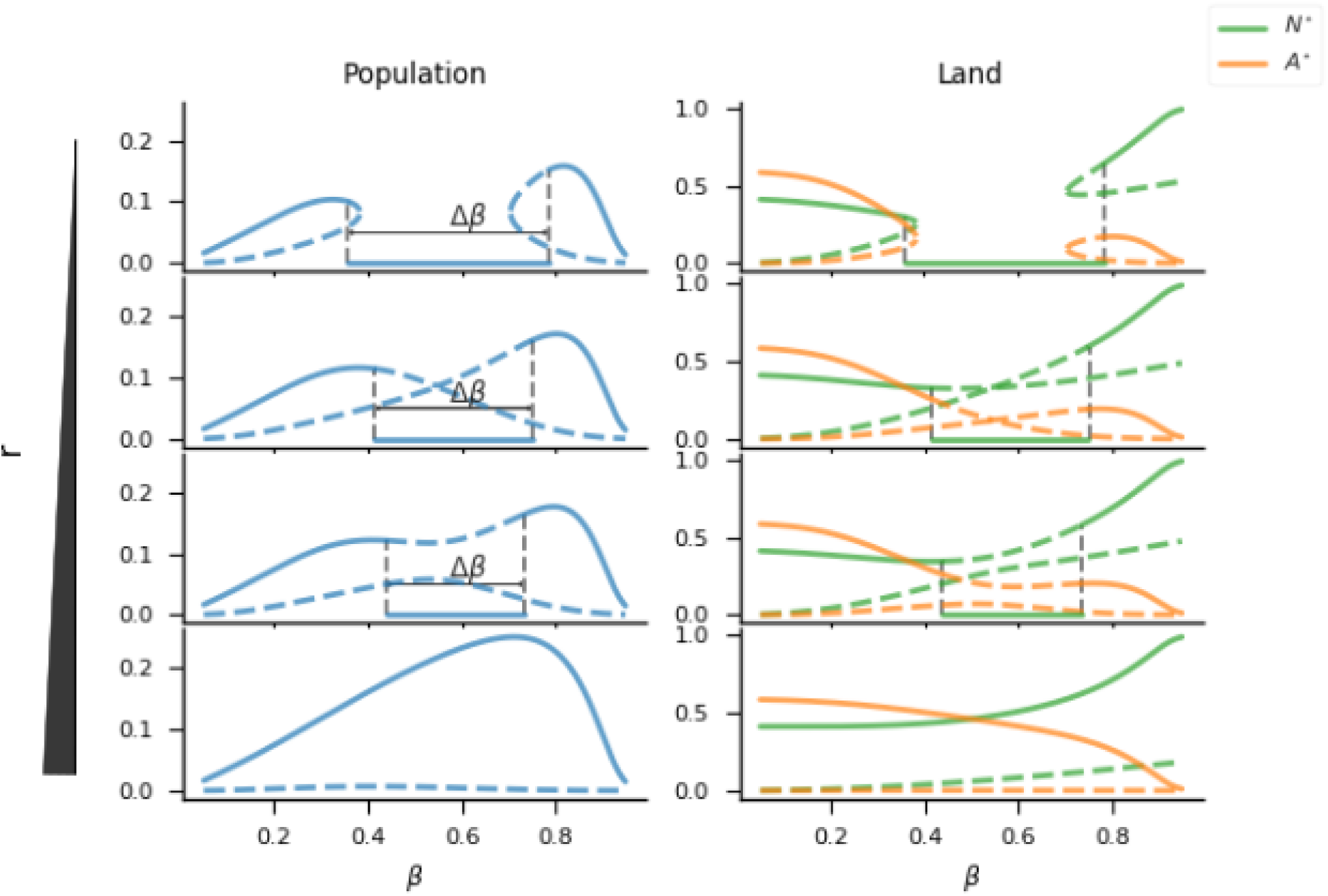
Bifurcation diagrams for the land use strategy parameter *β* and for different values of natural land’s recovery capacity. Socio-ecological steady states are plotted as a function of the land use strategy. The dotted lines correspond to the unstable equilibria, and the solid ones to the stable ones. The size of the collapse region is Δ*β*. From the top to the bottom, the natural land’s recovery capacity increases. The values of the recovery capacity increase from top to bottom and were chosen to give a full picture of the steady state branches’ behaviour. Δ*β* decreases until disappearance as the recovery rate *r* increases. Parameter values for these simulation are *r* = 0.9, 0.97139, 1, 2, *d* = 1.0, *k* = 4.5, *e* = 1.0, *k*_0_ = 0.5 and *q* = 1.0.

**Figure 7:**
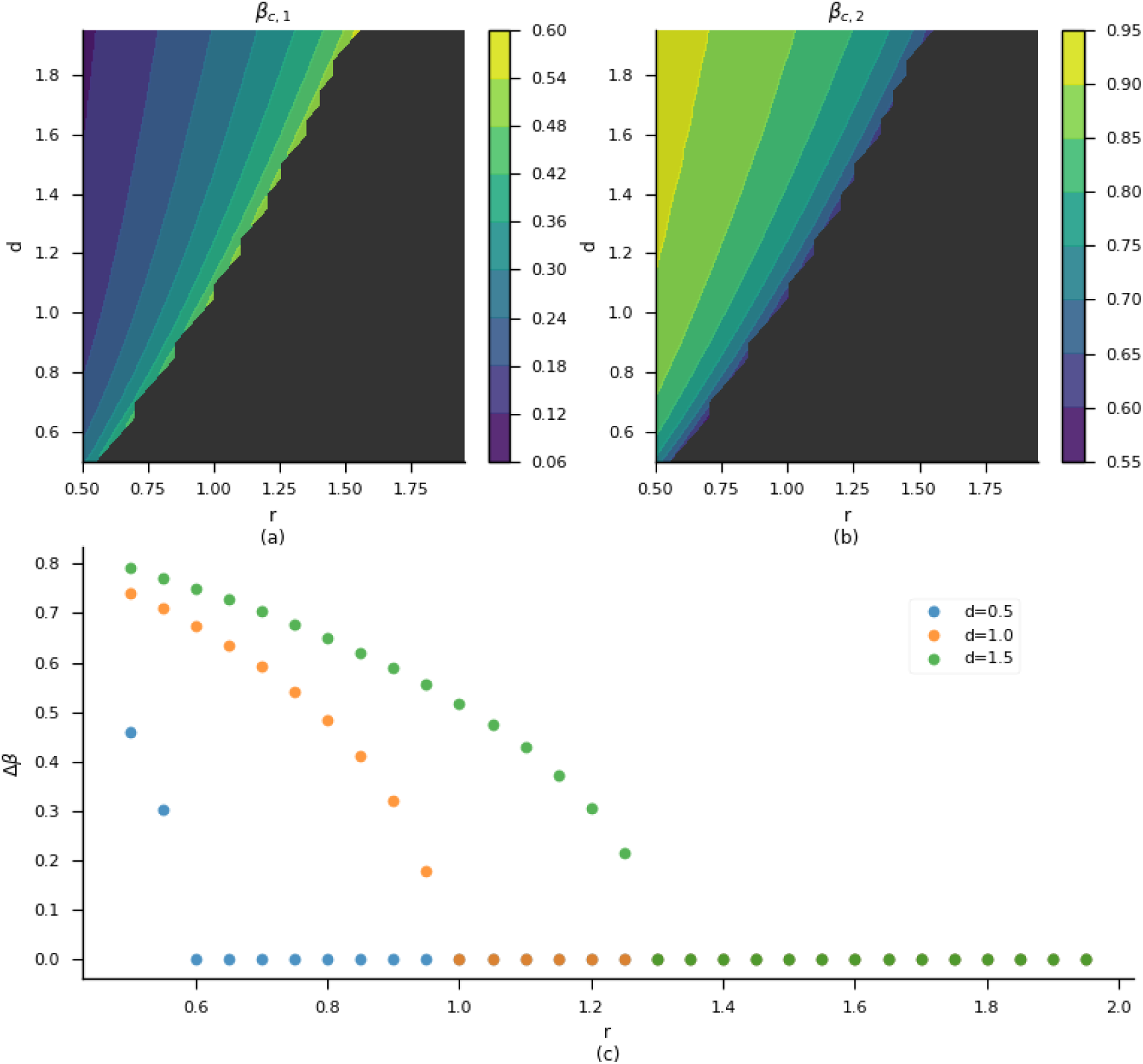
Collapse range size Δ*β* in function of landscape’s intrinsic characteristics. *On the top:* critical values *β*_*c*,1_ (a) and *β*_*c*,2_ (b) in function of the land’s recovery *r* and degradation *d* rates. The black colour depicts the region of the parameter space (*r, d*) where the human population is viable no matter the land use strategy. *On the bottom:* Size of the collapse range Δ*β* = *β*_*c*,1_ *– β*_*c*,2_ in function of the land’s recovery rate for different degradation rates. The amplitude of the collapse region sharply increases when *r* decreases. The parameter values are *k* = 4.5, *e* = 1.0, *k*_0_ = 0.5 and *q* = 1.0.

On the sparing side of the spectrum, when *β > β*_*c*,2_, the decrease in the land conversion effort that accompanies the increase of intensification succeeds in sparing natural land, and allows larger populations to exist in a landscape with a higher fraction of natural area. At the extreme of the sparing strategy spectrum, populations decrease when *β* rises. This is due to the decrease in the land conversion effort. Only a small fraction of land is converted to agriculture and as such a much lower population can be sustained with the same consumption level.

We investigated in more detail the relationship between the size of the critical region Δ*β* and the landscape intrinsic parameters *r* and *d* (Figure 7). The controur plots of Figure 7 show the variation of the critical values *β*_*c*,1_ and *β*_*c*,2_ as a function of *r* and *d*. For a given natural land degradation rate (*d*), increasing the recovery rate of degraded land (*r*) rises *β*_*c*,1_ and diminishes *β*_*c*,2_. When the difference between the two critical values reaches 0, the collapse range ceases to exist (black region in contour plots of Figure 7). The non-linearity of the edge between the coloured (Δ*β >* 0) and black regions (Δ*β* = 0) of the contour plots shows that the collapse frontier is more sensitive to *r* than *d*, such that when an increase in degradation requires a smaller increase in *r* to off-set the increase in degradation.

### 3.4 The dangers of naive agricultural land use planning

As it is formulated, our model does not allow us to know how much the land conversion effort should diminish for a given increase in agricultural intensification, in order to avoid socio-ecological collapse. This is because we fixed a linear trade-off between the conversion effort and the intensity. In reality, the relationship between them can be highly nonlinear. In order to address the question, we release the linear trade-off assumption, and let the land conversion effort to be independent of agricultural intensity. We then explore land use strategies along the two dimensions of intensification and extensification. In practice, this means we now have two parameters *K* (land conversion effort) and *β* (agricultural intensity) to describe a land use strategy instead of a single one. Hence the equations for land become:

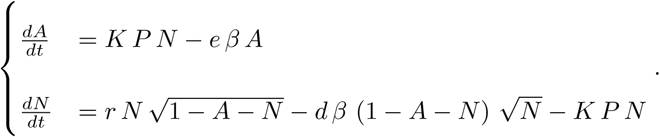

In Figure 8 we plot the regions of the land use strategy space, defined by the land conversion effort *K* and the agricultural intensity *β* where either the collapse equilibrium or the viable equilibrium are attained. The border between the two regions is fconcave rather than linear, which explains the existence of the collapse range Δ*β* we previously described. A linear decrease in the conversion effort in relation to agricultural intensity (solid black line in the graph) makes it unavoidable to cross the border between the viable and collapse equilibria. This result shows the non-triviality of designing sustainable land use strategies.

**Figure 8:**
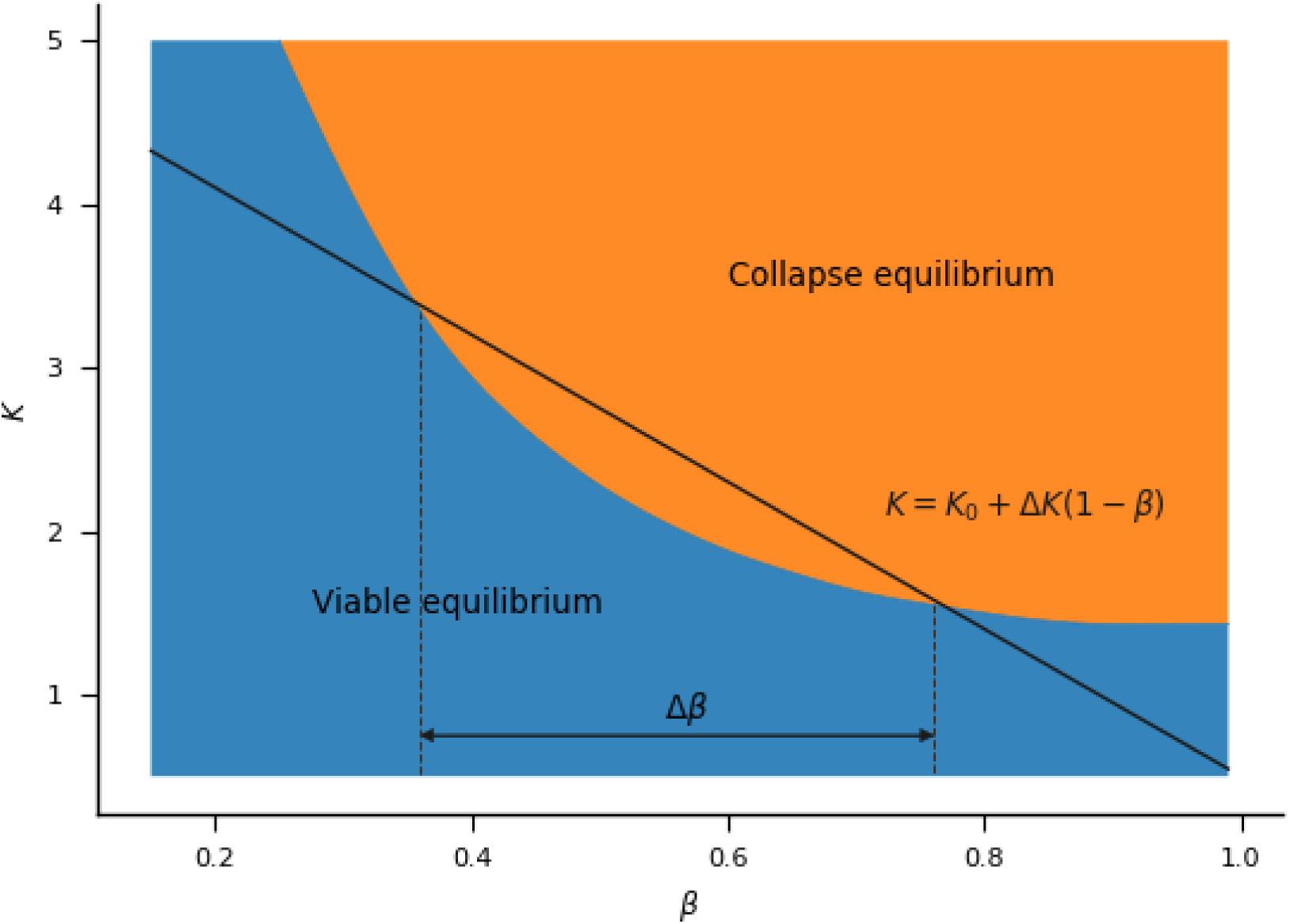
Long term system’s behaviour in the two-dimensional land use strategy space. Land use strategy is defined by the couple intensification *β* and conversion effort *K*. The blue region corresponds to the set of strategies leading to a viable socio-ecological equilibrium and the orange region corresponds to the ones leading to socio-ecological collapse. The solid black line depicts the linear trade-off between intensification and conversion effort that we were previously assuming. The border between the two regions is non-linear, which explains the existence of the collapse range Δ*β*.

## 4 Discussion

We investigated the impact of different land use strategies on the long-term sustainability of an agriculturally-based human society. Inspired by the land sparing-sharing debate we introduced in our model a land use strategy parameter (*β*) that controls the trade-off between agricultural intensity and land conversion effort. We then studied the behaviour of the coupled socio-ecological system across a continuum of strategies ranging from low agricultural intensity and high conversion effort (*β* = 0) to high agricultural intensity and low conversion effort (*β* = 1). We focused on characterising the set of land use strategies that lead to socio-ecological collapse as a function of the land’s recovery capacity. Our results show how agricultural intensification leads to irreversible land degradation and population collapse when not accompanied by a strong reduction of the land conversion effort. We highlighted the non-trivial relationship between an increase in agricultural intensification and the required decrease in the conversion effort. Naive land use planning can drive the socio-ecological system to a critical transition that undermines sustainability and leads to irreversible collapse.

Our model predicts that the most suitable strategy to ally a large population size and nature conservation is to practice extremely intense agriculture and minimise the conversion of natural land to agriculture. On the opposite side of the spectrum, low agricultural intensification and high conversion efforts lead to preserved landscapes but also to significantly lower population sizes. Therefore, for our current population the model seems to support the advocates of the sparing hypothesis. However, the existence of a collapse region in the middle of the strategy spectrum suggests that it is not simply a question of sparing.

For a certain amount of time, which depends on the speed of technological development, a nonviable land use strategy may be adopted. Therefore, if technological development in the agriculture sector stagnates there is a greater risk of getting trapped in the collapse region. We have not explored the effect of dynamic strategies in our model, but we are able to deduce from the model that strategies inside the collapse region push the landscape beneath a tipping point. Thus, if the land use strategy does not cross the collapse region fast enough, socio-ecological collapse will happen, even if a viable strategy is eventually adopted. Whether the levels of intensification required to overcome the collapse region are attainable or how to accurately measure intensity and know where we are on the spectrum are unclear.

We modelled the trade-off between expansion and intensification by imposing a negative linear relationship between agricultural intensity and conversion effort, thus reducing the two dimensions of the strategy to a single parameter along an axis that mimics the sparing-sharing one.

It is evident that simultaneous increases in both agricultural expansion and intensification cannot be viable in the long term. However, current practices favour both intensification and expansion. Since the adoption of agriculture by humans and the apparition of permanent settlements, agricultural intensification has fuelled urban growth. The last century’s Green Revolution is a recent and striking example – agricultural intensification increased yields and food security. However, this was also the period of fastest population growth in history, which further increased demand and motivated agricultural intensification and expansion. The societal and economic benefits of agricultural intensification that ensure food security are undeniable. However, agricultural intensification and expansion have caused several environmental problems such as soil erosion, nutrient runoff, water pollution or habitat destruction and fragmentation. It has also caused profound societal transformations, in particular the disappearance of small agricultural producers, that fuel urbanisation and change consumption patterns.

Agricultural intensification is considered a plausible explanation of past societal collapses, such as in the Roman or Mayan examples (Diamond, 2005). In a recent study, Roman et al. (2018) proposed a model that reproduced the Mayan population estimates and the model results indicated that agricultural intensification might have played a crucial role in the collapse. Without focusing on a particular historical case, our model also illustrates how agricultural intensification can lead to socio-ecological collapse when the conservation of natural lands is not large enough. This is extremely pertinent in the context of the current environmental crisis.

Population collapse emerges from our model as a consequence of large-scale land degradation, which impairs agricultural production in two ways: first, the deterioration of the landscape critically depletes the stock of natural land, which is the primary source for conversion to agriculture; second, natural land depletion reduces the provision of ecosystem services to existing agricultural areas. Since land degradation can critically decrease agricultural production, it seems ironic that agricultural land use is nowadays considered to be among the major causes of land degradation. This dangerous feedback loop poses a serious threat to sustainability as agricultural expansion and/or intensification to cope with reduced production can further reduce production in the long term by accentuating land degradation. Thus, we stress the major importance that agricultural land use planning has on the sustainability of socio-ecological systems.

Our results also stress the importance of careful land use planning to achieve sustainability. By releasing the linear trade-off between land conversion and agricultural intensity in our model, we showed that sustainable land use strategies can be obtained for the entire spectrum of intensification we consider. However, the frontier between unsustainable and sustainable land use can be far for trivial. The existence of unsustainable land use strategies comes from a bad evaluation of the needed reduction of land conversion for a given increase on intensification. As our model is a simplification of real population dynamics and land use planning, we do not claim that the frontier between sustainable and collapse paths is as we describe. However, we show that it is very likely for this frontier to be far from trivial, hence making it easy for naive land use planning to fail.

Since the beginning of agriculture, the fraction of the world’s agricultural land has steadily increased, and with it the fraction of natural land has decreased. Even if globally, at an aggregated scale, it could be argued that we have not yet reached a critical point or planetary boundary (Steffen et al., 2015), at a local scale this might not be true. Agricultural land use is not spatially homogeneous, and agricultural production is often strongly localised: the Pampas in South America, and the Great Plains in the United States are two examples. Moreover, these major agricultural regions, are mostly expansive, and intensive monocultures. Hence, at a regional scale there is neither sparing nor sharing, rather extensive exploitation, which makes the landscape highly susceptible to irreversible ecological degradation. In South American grasslands and forests, these practices have already caused major environmental degradation (Guerschman and Paruelo, 2005; Fearnside, 2001; Pengue, 2005), in addition to societal problems (Pengue, 2005). Agricultural expansion has already destroyed most of Brazil’s Atlantic Forest (Science, 2003; Ribeiro et al., 2011), and now it is advancing over the Amazon Forest, one of the world’s biodiversity hotspots (Soares-Filho et al., 2006; Davidson et al., 2012; Lovejoy and Nobre, 2018; Nepstad et al., 2008). The possibility of local collapses poses a threat to global sustainability, as it is unclear how these local collapses can propagate over the world, via environmental degradation but also via changes in trade or migration networks.

Environmental degradation in these major agricultural regions has not yet had an impact on human population via production drops. Quite the contrary, agricultural yields have increased over time. In the US, the potential negative consequences of agricultural production have been compensated by an increased use of fertilisers and pesticides over the six last decades (Wang, 2015). However, the growth rate of agricultural productivity in high and middle income countries, including the US and Europe, has been slowing down in recent years (Alston et al., 2010). This slowdown has not yet appeared in South America but it might just be a matter of time (Trindade and Fulginiti, 2015).

Our model is a simplified representation of agricultural practices and population growth. In particular, we have assumed that humans cannot adapt to environmental degradation by changing land use strategies or fertility or consumption behaviours over time. This is a major limitation, as adaptive strategies could potentially prevent the predicted population collapse. However, this does not reduce the pertinence of our results, as the current trends have the potential to cause a collapse, if habits go unchanged. Moreover, if humans succeed, through changes in cultural patterns, to avoid a drastic population reduction, it is very likely that the changes will be dramatic and involve, for example, a complete socio-economical restructuring, which can also be interpreted as a collapse (Cumming and Peterson, 2017). Hence, our results highlight that our current socio-ecological system might be heading towards dramatic changes, even though it is hard to predict the form they will take.

## 5 Conclusions

By exploring a continuum of strategies, our work differentiates from previous models of coupled human-land dynamics that studied the transition to higher agricultural intensification levels by considering two discrete levels of intensity (Roman et al., 2018). We modelled agricultural land use planning along two dimensions: expansion, given by the population’s conversion effort, and intensification. Expansion and intensification can act in synergy to increase landscape degradation, but there are also trade-offs between them. Agricultural expansion increases the stock of potential degraded land, while agricultural intensification can both speed and deepen agricultural land’s degradation.

Our results show how increasing agricultural intensification leads to socio-ecological collapse when there is an insufficient reduction of the land conversion effort. Agricultural intensification increases agricultural production, hence human population size, if consumption levels are kept equal. Population growth feeds back on the landscape’s composition by further accelerating land conversion. Eventually, land degradation reduces resource production and causes the population to overshoot its carrying capacity and ultimately decline.

The reduced human population size lowers the pressure on the natural areas and in some cases, this stabilises the system and makes it viable in the long term. However, when the per capita land conversion effort is large compared to the degree of intensification, the human population pushes the landscape beyond a tipping point after which there is not enough natural land to ensure recovery of the degraded land, making collapse inevitable.

By modelling the bi-directional feedbacks between human demography and land use, we have shown how misguided or naive agricultural land use planning can lead a socio-ecological system to collapse. Our model illustrates a potential mechanism that may explain the collapse of past societies but also a possible future collapse. As the gloabal human population is projected to keep growing in the next decades, adaptation through the reduction of consumption levels seems necessary to reduce the chance of a potential future collapse.

## Acknowledgements

We thank Matthieu Barbier for fruitful discussions. This work was supported by the TULIP Laboratory of Excellence (ANR-10-LABX-41) and was conducted within the framework of the BIOSTASES Advanced Grant funded by the European Research Council under the European Union’s Horizon 2020 research and innovation programme (grant agreement no. 666971).

## References

Teja Tscharntke, Yann Clough, Thomas C. Wanger, Louise Jackson, Iris Motzke, Ivette Perfecto, John Vandermeer, and Anthony Whitbread. Global food security, biodiversity conservation and the future of agricultural intensification. Biological Conservation, 151(1):53–59, July 2012. ISSN 0006-3207. doi:10.1016/j.biocon.2012.01.068.

John Bongaarts. IPBES, 2019. Summary for policymakers of the global assessment report on biodiversity and ecosystem services of the Intergovernmental Science-Policy Platform on Biodiversity and Ecosystem Services. Population and Development Review, 45(3):680–681, 2019. ISSN 1728-4457. doi: 10.1111/padr.12283.

C. Corvalán, Simon Hales, A. J. McMichael, Millennium Ecosystem Assessment (Program), and World Health Organization, editors. Ecosystems and Human Well-Being: Health Synthesis. Millennium Ecosystem Assessment. World Health Organization, Geneva, Switzerland, 2005. ISBN 978-92-4-156309-3. OCLC: ocm62587759.

Andrew P. Jacobson, Jason Riggio, Alexander M. Tait, and Jonathan E. M. Baillie. Global areas of low human impact (‘Low Impact Areas’) and fragmentation of the natural world. Sci Rep, 9(1):1–13, October 2019. ISSN 2045-2322. doi: 10.1038/s41598-019-50558-6.

J. Nowosad and T. F. Stepinski. Stochastic, Empirically Informed Model of Landscape Dynamics and Its Application to Deforestation Scenarios. Geophysical Research Letters, 46(23):13845–13852, 2019. ISSN 1944-8007. doi: 10.1029/2019GL085952.

Matthew G. E. Mitchell, Elena M. Bennett, and Andrew Gonzalez. Forest fragments modulate the provision of multiple ecosystem services. Journal of Applied Ecology, 51(4):909–918, August 2014. ISSN 00218901. doi: 10.1111/1365-2664.12241.

Alison G. Power. Ecosystem services and agriculture: Tradeoffs and synergies. Philosophical Transactions of the Royal Society B: Biological Sciences, 365(1554):2959–2971, September 2010. doi: 10.1098/rstb.2010.0143.

Matthew G. E. Mitchell, Elena M. Bennett, and Andrew Gonzalez. Linking Landscape Connectivity and Ecosystem Service Provision: Current Knowledge and Research Gaps. Ecosystems, 16(5):894–908, August 2013. ISSN 1432-9840, 1435-0629. doi: 10.1007/s10021-013-9647-2.

Victor Cazalis, Michel Loreau, and Kirsten Henderson. Do we have to choose between feeding the human population and conserving nature? Modelling the global dependence of people on ecosystem services. Science of The Total Environment, 634:1463–1474, September 2018. ISSN 0048-9697. doi: 10.1016/j.scitotenv.2018.03.360.

Leon C. Braat and Rudolf de Groot. The ecosystem services agenda:bridging the worlds of natural science and economics, conservation and development, and public and private policy. Ecosystem Services, 1(1): 4–15, July 2012. ISSN 2212-0416. doi: 10.1016/j.ecoser.2012.07.011.

Joseph Tainter. The Collapse of Complex Societies. Cambridge University Press, 1988. ISBN 978-0-521-38673-9.

Graeme S. Cumming and Garry D. Peterson. Unifying Research on Social–Ecological Resilience and Collapse. Trends in Ecology & Evolution, 32(9):695–713, September 2017. ISSN 0169-5347. doi: 10.1016/j.tree.2017.06.014.

Jared M. Diamond. Collapse: How Societies Choose to Fail or Succeed. New York: Viking, 2005.

James A. Brander and M. Scott Taylor. The Simple Economics of Easter Island: A Ricardo-Malthus Model of Renewable Resource Use. The American Economic Review, 88(1):119–138, 1998. ISSN 0002-8282.

Gunnar Brandt and Agostino Merico. The slow demise of Easter Island: Insights from a modeling investigation. Front. Ecol. Evol., 3, 2015. ISSN 2296-701X. doi: 10.3389/fevo.2015.00013.

Thomas R. Dalton and R. Morris Coats. Could institutional reform have saved Easter Island? J Evol Econ, 10(5):489–505, September 2000. ISSN 1432-1386. doi: 10.1007/s001910000050.

David H. Good and Rafael Reuveny. The fate of Easter Island: The limits of resource management institutions. Ecological Economics, 58(3):473–490, June 2006. ISSN 0921-8009. doi: 10.1016/j.ecolecon.2005.07.022.

John C. V. Pezzey and John M. Anderies. The effect of subsistence on collapse and institutional adaptation in population–resource societies. Journal of Development Economics, 72(1):299–320, October 2003. ISSN 0304-3878. doi: 10.1016/S0304-3878(03)00078-6.

Rafael Reuveny and Christopher S. Decker. Easter Island: Historical anecdote or warning for the future? Ecological Economics, 35(2):271–287, November 2000. ISSN 0921-8009. doi: 10.1016/S0921-8009(00)00202-0.

Robert L. Axtell, Joshua M. Epstein, Jeffrey S. Dean, George J. Gumerman, Alan C. Swedlund, Jason Harburger, Shubha Chakravarty, Ross Hammond, Jon Parker, and Miles Parker. Population growth and collapse in a multiagent model of the Kayenta Anasazi in Long House Valley. PNAS, 99(suppl 3):7275–7279, May 2002. ISSN 0027-8424, 1091-6490. doi: 10.1073/pnas.092080799.

Timothy A. Kohler, R. Kyle Bocinsky, Denton Cockburn, Stefani A. Crabtree, Mark D. Varien, Kenneth E. Kolm, Schaun Smith, Scott G. Ortman, and Ziad Kobti. Modelling prehispanic Pueblo societies in their ecosystems. Ecological Modelling, 241:30–41, August 2012. ISSN 0304-3800. doi: 10.1016/j.ecolmodel.2012.01.002.

Scott Heckbert. MayaSim: An Agent-Based Model of the Ancient Maya Social-Ecological System. JASSS, 16(4):11, 2013. ISSN 1460-7425.

Linda Kuil, Gemma Carr, Alberto Viglione, Alexia Prskawetz, and Günter Blöschl. Conceptualizing socio-hydrological drought processes: The case of the Maya collapse. Water Resources Research, 52(8):6222–6242, 2016. ISSN 1944-7973. doi: 10.1002/2015WR018298.

Sabin Roman, Erika Palmer, and Markus Brede. The Dynamics of Human–Environment Interactions in the Collapse of the Classic Maya. Ecological Economics, 146:312–324, April 2018. ISSN 0921-8009. doi: 10.1016/j.ecolecon.2017.11.007.

Scott Barrett, Aisha Dasgupta, Partha Dasgupta, W. Neil Adger, John Anderies, Jeroen van den Bergh, Caroline Bledsoe, John Bongaarts, Stephen Carpenter, F. Stuart Chapin, Anne-Sophie Crépin, Gretchen Daily, Paul Ehrlich, Carl Folke, Nils Kautsky, Eric F. Lambin, Simon A. Levin, Karl-Göran Mäler, Rosamond Naylor, Karine Nyborg, Stephen Polasky, Marten Scheffer, Jason Shogren, Peter Søgaard Jørgensen, Brian Walker, and James Wilen. Social dimensions of fertility behavior and consumption patterns in the Anthropocene. Proceedings of the National Academy of Sciences, 117(12):6300–6307, March 2020. ISSN 0027-8424, 1091-6490. doi: 10.1073/pnas.1909857117.

Safa Motesharrei, Jorge Rivas, and Eugenia Kalnay. Human and nature dynamics (HANDY): Modeling inequality and use of resources in the collapse or sustainability of societies. Ecological Economics, 101: 90–102, May 2014. ISSN 0921-8009. doi: 10.1016/j.ecolecon.2014.02.014.

Kirsten Henderson and Michel Loreau. How ecological feedbacks between human population and land cover influence sustainability. PLOS Computational Biology, 14(8):e1006389, August 2018. ISSN 1553-7358. doi: 10.1371/journal.pcbi.1006389.

Kirsten Henderson and Michel Loreau. An ecological theory of changing human population dynamics. People and Nature, 1(1):31–43, 2019. ISSN 2575-8314. doi: 10.1002/pan3.8.

A. S. Lafuite and M. Loreau. Time-delayed biodiversity feedbacks and the sustainability of social-ecological systems. Ecological Modelling, 351:96–108, May 2017. ISSN 0304-3800. doi: 10.1016/j.ecolmodel.2017.02.022.

A.-S. Lafuite, C. de Mazancourt, and M. Loreau. Delayed behavioural shifts undermine the sustainability of social–ecological systems. Proceedings of the Royal Society B: Biological Sciences, 284(1868):20171192, December 2017. ISSN 0962-8452, 1471-2954. doi: 10.1098/rspb.2017.1192.

A. S. Lafuite, G. Denise, and M. Loreau. Sustainable Land-use Management Under Biodiversity Lag Effects. Ecological Economics, 154:272–281, December 2018. ISSN 0921-8009. doi: 10.1016/j.ecolecon.2018.08.003.

Sean S. Downey, W. Randall Haas, and Stephen J. Shennan. European Neolithic societies showed early warning signals of population collapse. Proceedings of the National Academy of Sciences, 113(35):9751–9756, August 2016. ISSN 0027-8424, 1091-6490. doi: 10.1073/pnas.1602504113.

Ian Kuijt and Nigel Goring-Morris. Foraging, Farming, and Social Complexity in the Pre-Pottery Neolithic of the Southern Levant: A Review and Synthesis. Journal of World Prehistory, 16(4):361–440, December 2002. ISSN 1573-7802. doi: 10.1023/A:1022973114090.

Ruth S. DeFries, Thomas Rudel, Maria Uriarte, and Matthew Hansen. Deforestation driven by urban population growth and agricultural trade in the twenty-first century. Nature Geosci, 3(3):178–181, March 2010. ISSN 1752-0894, 1752-0908. doi: 10.1038/ngeo756.

Amy Lilienfeld and Mette Asmild. Estimation of excess water use in irrigated agriculture: A Data Envelopment Analysis approach. Agricultural Water Management, 94(1):73–82, December 2007. ISSN 0378-3774. doi: 10.1016/j.agwat.2007.08.005.

Maria A. Tsiafouli, Elisa Thébault, Stefanos P. Sgardelis, Peter C. de Ruiter, Wim H. van der Putten, Klaus Birkhofer, Lia Hemerik, Franciska T. de Vries, Richard D. Bardgett, Mark Vincent Brady, Lisa Bjornlund, Helene Bracht Jørgensen, Sören Christensen, Tina D’ Hertefeldt, Stefan Hotes, W. H. Gera Hol, Jan Frouz, Mira Liiri, Simon R. Mortimer, Heikki Setälä, Joseph Tzanopoulos, Karoline Uteseny, Václav Pižl, Josef Stary, Volkmar Wolters, and Katarina Hedlund. Intensive agriculture reduces soil biodiversity across Europe. Global Change Biology, 21(2):973–985, 2015. ISSN 1365-2486. doi: 10.1111/gcb.12752.

John N. Quinton, Gerard Govers, Kristof Van Oost, and Richard D. Bardgett. The impact of agricultural soil erosion on biogeochemical cycling. Nature Geosci, 3(5):311–314, May 2010. ISSN 1752-0908. doi: 10.1038/ngeo838.

Eric A. Davidson, Alessandro C. de Araújo, Paulo Artaxo, Jennifer K. Balch, I. Foster Brown, Mercedes M. C. Bustamante, Michael T. Coe, Ruth S. DeFries, Michael Keller, Marcos Longo, J. William Munger, Wilfrid Schroeder, Britaldo S. Soares-Filho, Carlos M. Souza, and Steven C. Wofsy. The Amazon basin in transition. Nature, 481(7381):321–328, January 2012. ISSN 1476-4687. doi: 10.1038/nature10717.

Ricardo Grau, Tobias Kuemmerle, and Leandro Macchi. Beyond ‘land sparing versus land sharing’: Environmental heterogeneity, globalization and the balance between agricultural production and nature conservation. Current Opinion in Environmental Sustainability, 5(5):477–483, October 2013. ISSN 18773435. doi: 10.1016/j.cosust.2013.06.001.

Ben Phalan, Malvika Onial, Andrew Balmford, and Rhys E. Green. Reconciling Food Production and Biodiversity Conservation: Land Sharing and Land Sparing Compared. Science, 333(6047):1289–1291, September 2011a. ISSN 0036-8075, 1095-9203. doi: 10.1126/science.1208742.

Ben Phalan, Andrew Balmford, Rhys E. Green, and Jörn P.W. Scharlemann. Minimising the harm to biodiversity of producing more food globally. Food Policy, 36:S62–S71, January 2011b. ISSN 03069192. doi: 10.1016/j.foodpol.2010.11.008.

Ben Balmford, Rhys E. Green, Malvika Onial, Ben Phalan, and Andrew Balmford. How imperfect can land sparing be before land sharing is more favourable for wild species? Journal of Applied Ecology, 56(1): 73–84, January 2019. ISSN 00218901. doi: 10.1111/1365-2664.13282.

I. Perfecto and J. Vandermeer. The agroecological matrix as alternative to the land-sparing/agriculture intensification model. Proceedings of the National Academy of Sciences, 107(13):5786–5791, March 2010. ISSN 0027-8424, 1091-6490. doi: 10.1073/pnas.0905455107.

Benjamin T. Phalan. What Have We Learned from the Land Sparing-sharing Model? Sustainability, 10(6):1760, June 2018. doi: 10.3390/su10061760.

Philip M. Fearnside. Soybean cultivation as a threat to the environment in Brazil. Environmental Conservation, 28(1):23–38, March 2001. ISSN 1469-4387, 0376-8929. doi: 10.1017/S0376892901000030.

Walter A. Pengue. Transgenic Crops in Argentina: The Ecological and Social Debt. Bulletin of Science, Technology & Society, 25(4):314–322, August 2005. ISSN 0270-4676. doi: 10.1177/0270467605277290.

Carlos Reboratti. Un mar de soja: La nueva agricultura en Argentina y sus consecuencias. Revista de geografía Norte Grande, (45):63–76, May 2010. ISSN 0718-3402. doi: 10.4067/S0718-34022010000100005.

Britaldo Silveira Soares-Filho, Daniel Curtis Nepstad, Lisa M. Curran, Gustavo Coutinho Cerqueira, Ricardo Alexandrino Garcia, Claudia Azevedo Ramos, Eliane Voll, Alice McDonald, Paul Lefebvre, and Peter Schlesinger. Modelling conservation in the Amazon basin. Nature, 440(7083):520–523, March 2006. ISSN 1476-4687. doi: 10.1038/nature04389.

Viki A. Cramer, Richard J. Hobbs, and Rachel J. Standish. What’s new about old fields? Land abandonment and ecosystem assembly. Trends in Ecology & Evolution, 23(2):104–112, February 2008. ISSN 0169-5347. doi: 10.1016/j.tree.2007.10.005.

Lander Baeten, Darline Velghe, Margot Vanhellemont, Pieter De Frenne, Martin Hermy, and Kris Verheyen. Early Trajectories of Spontaneous Vegetation Recovery after Intensive Agricultural Land Use. Restoration Ecology, 18:379–386, March 2010. ISSN 1526-100X. doi: 10.1111/j.1526-100X.2009.00627.x.

Yuval R. Zelnik, Hannes Uecker, Ulrike Feudel, and Ehud Meron. Desertification by front propagation? Journal of Theoretical Biology, 418:27–35, April 2017. ISSN 0022-5193. doi: 10.1016/j.jtbi.2017.01.029.

Yuval R. Zelnik and Ehud Meron. Regime shifts by front dynamics. Ecological Indicators, 94:544–552, November 2018. ISSN 1470-160X. doi: 10.1016/j.ecolind.2017.10.068.

Samir Suweis, Andrea Rinaldo, Amos Maritan, and Paolo D’Odorico. Water-controlled wealth of nations. Proceedings of the National Academy of Sciences, 110(11):4230–4233, March 2013. ISSN 0027-8424, 1091-6490. doi: 10.1073/pnas.1222452110.

Amy Goldberg, Alexis M. Mychajliw, and Elizabeth A. Hadly. Post-invasion demography of prehistoric humans in South America. Nature, 532(7598):232–235, April 2016. ISSN 1476-4687. doi: 10.1038/nature17176.

Matthew G. E. Mitchell, Andrés F. Suarez-Castro, Maria Martinez-Harms, Martine Maron, Clive McAlpine, Kevin J. Gaston, Kasper Johansen, and Jonathan R. Rhodes. Reframing landscape fragmentation’s effects on ecosystem services. Trends in Ecology & Evolution, 30(4):190–198, April 2015. ISSN 0169-5347. doi: 10.1016/j.tree.2015.01.011.

Will Steffen, Katherine Richardson, Johan Rockström, Sarah E. Cornell, Ingo Fetzer, Elena M. Bennett, Reinette Biggs, Stephen R. Carpenter, Wim de Vries, Cynthia A. de Wit, Carl Folke, Dieter Gerten, Jens Heinke, Georgina M. Mace, Linn M. Persson, Veerabhadran Ramanathan, Belinda Reyers, and Sverker Sörlin. Planetary boundaries: Guiding human development on a changing planet. Science, 347(6223), February 2015. ISSN 0036-8075, 1095-9203. doi: 10.1126/science.1259855.

Juan Pablo Guerschman and José María Paruelo. Agricultural impacts on ecosystem functioning in temperate areas of North and South America. Global and Planetary Change, 47(2):170–180, July 2005. ISSN 0921-8181. doi: 10.1016/j.gloplacha.2004.10.021.

Centre for Applied Biodiversity Science. The Atlantic Forest of South America: Biodiversity Status, Threats, and Outlook. Island Press, 2003. ISBN 978-1-55963-989-7.

Milton Cezar Ribeiro, Alexandre Camargo Martensen, Jean Paul Metzger, Marcelo Tabarelli, Fábio Scarano, and Marie-Josee Fortin. The Brazilian Atlantic Forest: A Shrinking Biodiversity Hotspot. In Frank E. Zachos and Jan Christian Habel, editors, Biodiversity Hotspots: Distribution and Protection of Conservation Priority Areas, pages 405–434. Springer, Berlin, Heidelberg, 2011. ISBN 978-3-642-20992-5. doi: 10.1007/978-3-642-20992-521.

Thomas E. Lovejoy and Carlos Nobre. Amazon Tipping Point. Science Advances, 4(2):eaat2340, February 2018. ISSN 2375-2548. doi: 10.1126/sciadv.aat2340.

Daniel C Nepstad, Claudia M Stickler, Britaldo Soares Filho, and Frank Merry. Interactions among Amazon land use, forests and climate: Prospects for a near-term forest tipping point. Philosophical Transactions of the Royal Society B: Biological Sciences, 363(1498):1737–1746, May 2008. doi: 10.1098/rstb.2007.0036.

Sun Ling Wang. Agricultural Productivity Growth in the United States: Measurement, Trends, and Drivers. page 78, 2015.

Julian Alston, Bruce Babcock, and Philip Pardey. The Shifting Patterns of Agricultural Production and Productivity Worldwide. CARD Books, January 2010.

Federico José Trindade and Lilyan Estela Fulginiti. Is there a slowdown in agricultural productivity growth in South America? Agricultural Economics, 46(S1):69–81, 2015. ISSN 1574-0862. doi: 10.1111/agec.12199.

